# Dissecting genetic correlation and pleiotropy through a genetic cross

**DOI:** 10.1101/2023.05.12.540583

**Authors:** Haoran Cai, Kerry Geiler-Samerotte, David L. Des Marais

## Abstract

Genetic correlation represents an important class of evolutionary constraint, which is itself evolvable. Empirical studies have found mixed results on whether genetic correlations change rapidly or slowly. This uncertainty challenges our ability to predict the outcome of selection. Despite the tremendous diversity and complexity of life forms, there are certain forms of life that are never observed. This might be because of developmental biases that restrict how organisms can evolve, or because they have low fitness in any environment yet available on Earth. Given that both developmental bias and selection can generate similar phenotypes, it is difficult to distinguish between the two causes of evolutionary stasis among related taxa. For example, remarkably invariant traits are observed spanning million years, such as wing shape in *Drosophila* wherein qualitative differences are rare within genera. Here, we ask whether the absence of certain combinations of traits, as indicated by genetic correlation, reflects developmental bias. However, much confusion and controversy remain over definitions of developmental bias, and probing it is challenging. We thus present a novel approach aiming to dissect genetic correlations and estimate the relative contribution of developmental bias in maintaining genetic correlations. We do so by leveraging a common but under-utilized type of data: genetic crosses. Through empirical analyses, we find that our approach can distinguish whether genetically correlated traits are developmentally constrained to covary. We also find that our developmental bias metric is an indicator of genetic correlation stability across conditions. Our framework presents a feasible way to dissect the mechanisms underlying genetic correlation and pleiotropy.

## Introduction

Genetic correlation represents an important class of evolutionary constraints (Maynard Smith *et al*., 1985; Clark, 1987), which affects future evolutionary trajectories. Yet, genetic correlations are themselves evolvable (Doroszuk *et al*., 2008; Dugand *et al*., 2021; Delph *et al*., 2011; Conner, 2002; Uller *et al*., 2018; Wagner & Altenberg, 1996; Rohner & Berger, 2023; Wagner *et al*., 2007) and maintained by both the selection of trait combinations and, in some cases, developmental bias (Dugand *et al*., 2021; Arnold, 1992). Natural selection may favor certain combinations of traits and thereby actively maintaining genetic correlation via pleiotropy or linkage disequilibruim. Such genetic correlation may then further inhibit traits from evolving independently towards a theoretical phenotypic optimum (Schluter, 1996). On the other hand, genetic correlation can be maintained by bias due to intrinsic attributes of the organism, by energy, or by the laws of physics, relative to the assumption of isotropic variation. This latter concept has been decribed as developmental constraint or developmental bias (Maynard Smith *et al*., 1985; Arnold, 1992; Cheverud, 1984; Rohner & Berger, 2023). Developmental bias represents the observation that perturbations to biological systems, such as mutation or environmental variation, will tend to produce some phenotypic variants more readily (Uller *et al*., 2018; Waddington, 1957). In spite of the numerous studies that address genetic correlation as an evolutionary constraint, much confusion and controversy remains over definitions and mechanism(s) causing genetic correlation, and the relative importance of different mechanisms in shaping evolutionary trajectories (Muir *et al*., 2023; Conner *et al*., 2011).

The theoretical underpinnings for genetic covariance as an evolutionary constraint are well-developed (Lande, 1979; Lande & Arnold, 1983; Walsh & Blows, 2009). Genetic covariance specifically describes trait covariance due to pleiotropic alleles, where a single locus has effects on two traits, or due to linkage disequilibrium of two loci (Lande, 1980; Lynch *et al*., 1998; Falconer *et al*., 1996; Conner *et al*., 2004). The genetic information summarized by genetic covariance is connected to evolutionary processes in complex ways. For example, evolution toward a phenotypic optimum for two traits may be restricted if selection favors two traits antagonistically but the traits are positively correlated. That is, adaptive evolution can be limited if the joint vector of selection is antagonistic to the trait correlations (Schluter, 1996; Lande & Arnold, 1983; Cai & Des Marais, 2023a; Walsh & Blows, 2009; Henry & Stinch-combe, 2023b). In some cases, such evolutionary constraint may persist over long time scales (McGlothlin *et al*., 2018; Opedal *et al*., 2023).

Straightforward applications of evolutionary quantitative genetic theory regarding the joint evolution of a pair of traits generally assume an invariant genetic covariance structure (**G**) over the time frame of interest. However, the stability of genetic covariances and how they evolve remain unclear and contentious (Turelli, 1988; Bürger & Lande, 1994; Arnold *et al*., 2008; Steppan *et al*., 2002; Milocco & Salazar-Ciudad, 2022; Loeschcke, 1987; Barton & Turelli, 1989). Empirical studies of the evolution of genetic covariance structure have found mixed results on whether genetic covariance changes rapidly or slowly. Some comparisons of **G** matrices between natural populations found no evidence of change in **G** (Delahaie *et al*., 2017; Arnold *et al*., 2008; Hangartner *et al*., 2020; Henry & Stinchcombe, 2023a), while others have found changes in genetic covariance in only a few generations, across populations, in response to selection, or across environmental conditions (Chakrabarty & Schielzeth, 2020; Milocco & Salazar-Ciudad, 2022; Eroukhmanoff & Svensson, 2011; Walter *et al*., 2018; Wood & Brodie III, 2015; Cai & Des Marais, 2023b; Henry & Stinchcombe, 2023a; Hudson *et al*., 2022; Scoville *et al*., 2009; Monroe *et al*., 2021). Generally, it is largely unknown what determines the stability of genetic covariance (Wood & Brodie III, 2015) and this uncertainty challenges our ability to predict the outcome of selection.

The persistence of correlational constraint and whether genetic correlation is a good predictor of long-term evolutionary divergence ultimately hinge on the underlying mechanism (s) that causes genetic correlation (Loeschcke, 1987; Conner *et al*., 2011, 2004). For example, genetic correlation due to pleiotropy or tight linkage are much more likely to cause evolutionary constraint than those caused by linkage disequilibrium between loosely linked loci (Conner *et al*., 2011, 2004; Conner, 2002). Correlations due to pleiotropy or tight linkage may persist in the absence of selection, while correlations caused by linkage disequilibrium between unlinked loci likely change when selection pressure change (Conner, 2002; Conner *et al*., 2004, 2011). Theoretical work has suggested that mutation-induced genetic correlation is more stable than selection-induced correlation (Jones *et al*., 2003). Insight into the role of mutational correlation may reveal why genetic correlations between some traits are more constant over a long period as compared to other pairs of traits and why, in some cases, genetic constraints can be readily degraded by natural or artificial selection. However, empirically discriminating between mutation and development from other mechanisms of genetic correlation is challenging.

One traditional approach to achieve this is through mutation accumulation (MA) lines to assess the spectrum of phenotypic variation generated by *de novo* mutation in the absence of drift or selection (Zalts & Yanai, 2017; Braendle *et al*., 2010; Houle *et al*., 2017). However, mutation accumulation experiment captures not only the propensity of the developmental system to vary but also heterogeneity in mutational rates across genome, and bias of mutation spectrum, which constrains the mutational space (Stoltzfus & McCandlish 2017; Rohner & Berger 2023; James *et al*. 2023; Agashe *et al*. 2023, e.g., more mutable single nucleotide, etc.). In addition to that, it provides a very limited view of the distribution of mutational effects because the number of loci and spontaneous mutations sampled in each study is low (Hodgins-Davis *et al*., 2019), not to mention the high costs of mutation accumulation experiments.

The phenotypic distribution of F2 cross has been previously used to infer what would be expected by chance under neutral evolution (Fraser, 2020), The underlying rationale is that the F2 phenotypes result from random combinations of the segregating alleles, which mimics the random directionality expected under neutral evolution. Here, we sought to assess the contribution of developmental bias (a neutral expectation) against natural selection in producing genetic correlation by using genetic crosses. We use the term horizontal pleiotropy (HP) to describe a locus has an effect on two traits, where such pleiotropic effect deviates from the effects of the other loci across a genome (Fig. 1 b,d). Conversely, developmental bias between traits generates the observation of consistent pleiotropic effect of loci throughout the genome on a given trait pair (Fig. 1 a,c). We use this consistency of pleiotropic effect throughout the genome to indicate developmental bias, *r*_*D*_. We reason that if two traits are correlated because of developmental bias, these two traits should be correlated regardless of which specific variant causes the effect.

**Figure 1:**
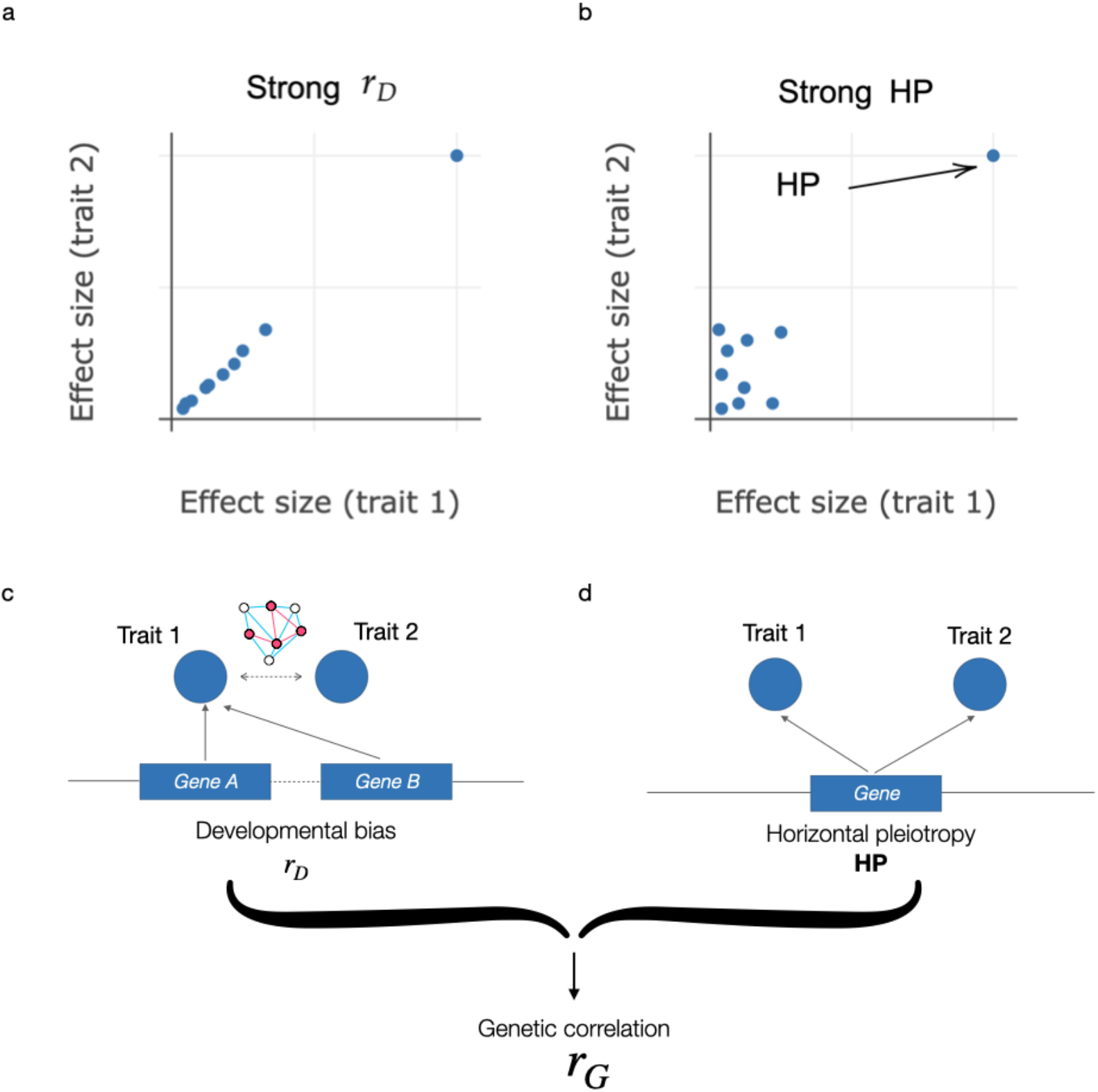
Conceptual framework for distinguishing between developmental bias and horizontal pleiotropy as drivers of genetic correlation (*r*_*G*_) between two traits. **a**. and **b**. Bi-plots showing the correlation of effect sizes of ten genetic loci on hypothetical traits 1 and 2. **a**. Strong developmental bias and low horizontal pleiotropy, as seen by coherent and consistent pleiotropic effect on the two traits across the sampled loci. **b**. One large-effect pleiotropic locus appears to drive the genetic correlation between traits 1 and 2, showing a strong horizontal pleiotropic (HP) effect. **c**. and **d**. Suggest genetic mechanisms for the observed effect correlations in **a** and **b. c**. A developmental bias *r*_*D*_, where each locus that affects one trait will inevitably affect the other trait, suggesting that the traits are inherently correlated regardless of the type and directions of genetic perturbation. **d**. Horizontal pleiotropy (HP), where a locus can have a direct effect on the two traits. A third cause of genetic correlation is linkage disequilibrium (LD; See Appendix B) where two loci, each determining a separate trait, exhibit nonrandom association of alleles between loci within populations.

Our primary goal in the present work is to dissect genetic correlations to understand underlying mechanisms maintaining the presence of genetic correlation. We do so by using both numerical simulations and data from a recombinant genetic mapping population. One key outcome is that we elaborate the definition of horizontal pleiotropy and identify QTLs that demonstrate high degree of horizontal pleiotropy. While another recent method exists for doing so (Geiler-Samerotte *et al*., 2020), our method is unique in that it does not require one to study clonal cells and can therefore be applied to a broader range of organisms. An additional goal of our study is to test our proposition that a genetic correlation that arises principally from genome-wide consistent pleiotropy is more persistent across conditions. When *r*_*G*_ are driven by numerious small effect size loci, we expect them to be more likely driven by developmental bias, as opposed to when they are driven by individual loci with large effect size horizontal pleiotropy, which are more likely maintained by present selection (McGuigan *et al*., 2014; Sella & Barton, 2019). In the latter case, any changes or perturbations affecting those specific loci (e.g., various types of environmental perturbations with QTL-by-environment effect, allele frequency changes etc.) may easily alter the genetic correlations. We find evidence that, indeed, the consistency of pleiotropy is an indicator of genetic correlation stability. We also show that genetic correlations driven by genome-wide consistent pleiotropy are likely to exhibit a highly polygenic architecture. Hence, a genetic correlation with a highly polygenic architecture may be more stable, corroborating previous expectations(Barton & Turelli, 1987, 1989; Lande, 1979). In sum, we use readily accessible QTL mapping data to understand how genetic architecture may be used to infer mechanisms of genetic correlation, to identify loci that act via high degree of horizontal pleiotropy, and to make predictions about how genetic correlations will change. These results suggest that this type of common data is under-utilized, and that analyzing recombinant populations with our approach can help to deepen our understandings of genetic correlation and pleiotropy.

## Results

During a genetic association study, each genetic marker is assigned an odds likelihood ratio along with an effect size for the trait of interest. Instead of identifying statistically significant loci in such conventional genetic association studies, the essential idea, here, is to examine the consistency of pleiotropic effects across genetic backgrounds. We define a locus with effects that deviate from the overall bivariate trend throughout the genome as a horizontal pleiotropic (HP) locus (Fig. 1b). We diagnose *r*_*D*_ as the consistency of pleiotropy across genetic backgrounds excluding HP loci. In a mapping population, allele substitutions at each locus represent non-directed (i.e., random) perturbations of varying directions and magnitudes. The additive effect of many loci thus are considered as random perturbations to an organism. To better illustrate the framework we propose, the bivariate effect size distributions under two scenarios are shown (Fig. 1a, b). The locus with a major phenotypic effect that deviates from the overall trend of other loci throughout the genome is an HP locus. Conversely, the consistency of pleiotropy (i.e., correlation among loci or locus-level correlation) is denoted as developmental bias *r*_*D*_ and the associated loci act via vertical pleiotropy.

### Simulation demonstrating relationships between consistency of pleiotropy and genetic correlation

To examine how the *r*_*D*_ – characterized by the effect size correlation among loci – relates to genetic correlation (*r*_*G*_), we first simulated two thousand trait pairs for a given simulated population with 500 individual genotypes. For each trait pair, genetic architecture with 226 loci was generated, with additive effect sizes sampled from a multivariate Laplace distribution.

The genetic values are obtained by multiplying the genotypes with allelic effect sizes, assuming no epistasis and no linkage disequilibruim. *r*_*G*_ is calculated by correlating the genetic values between two traits following standard protocols (Falconer *et al*., 1996). We calculated *r*_*D*_ and corresponding *r*_*G*_ for each pair of traits. Notably, *r*_*G*_ is a correlation across a population of individuals while *r*_*D*_ is a locus level correlation across a population of loci in a genome.

Assuming that there is no horizontal pleiotropy (HP) and no linkage disequilibrium (LD), we expect the effect size correlation and *r*_*G*_ to be equal. As expected, without accounting for HP, effect size correlation strongly correlates with *r*_*G*_ regardless of the genetic regimes (Fig. 2b,c). Next, we repeated our simulations under conditions with HP to understand how these forces would affect the correlations (Fig. 2d,e). Under the HP scenario, *n* randomly selected SNPs (0 *< n <* 10) are forced to have an HP effect, either concordant to or antagonistic with the rest of the loci. The genetic correlation *r*_*G*_ is not perfectly correlated with *r*_*D*_ under scenarios with HP, especially when the kurtosis of the effect size distribution is high. In an extreme case, a single large-effect locus can drive the genetic correlation despite the low *r*_*D*_ (Fig. S1). Collectively, these observations suggest that with HP and low polygenicity, *r*_*D*_ can vary in a wide range despite that *r*_*G*_ of these pairs are the same. With the pervasive empirical observations of horizontal pleiotropy (Verbanck *et al*., 2018; Jordan *et al*., 2019), we believe that what we proposed as developmental bias *r*_*D*_ is a distinguishable measure from *r*_*G*_.

**Figure 2:**
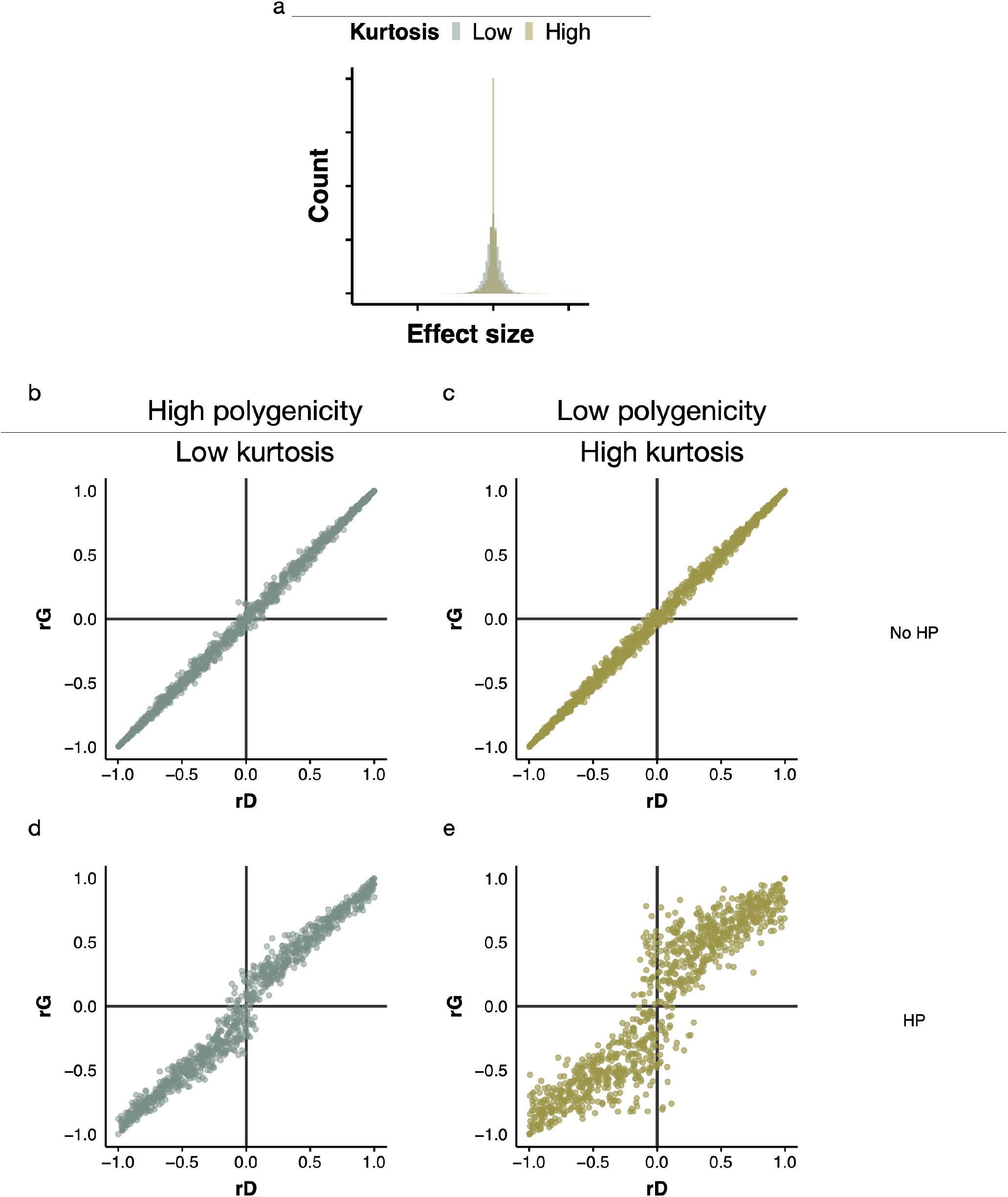
Simulations showing the relationship between developmental bias, *r*_*D*_, and genetic correlation, *r*_*G*_. We simulated 2,000 pairs of traits using an exponential model, with varying *σ* among the plots shown. *σ* describes the correlation of effect size when generating effect sizes for each trait pair. A second parameter of the exponential model, *γ*, indicates regimes of varying kurtosis of the effect size distribution: large *γ* represents low kurtosis while smaller *γ* represents a regime with high kurtosis, as shown in **a**.. Two regimes of genetic architecture are considered, with *γ* equals 1.0 (plots **b**. and **d**.) and 0.5 (plots **c**. and **e**.) Two scenarios are simulated. **b**. and **c**. Under the first scenario, no horizontal pleiotropy (HP), the pattern does not change with the kurtosis and *r*_*G*_ nearly perfectly represents *r*_*D*_. **d**. and **e**. Under the second scenario, to introduce HP, 0 to 10 randomly chosen SNP are introduced for each trait pair, with a shared pleiotropic effect for two traits, regardless of the pleiotropic effect of remaining loci.

Our additional individual-based simulation using NEMO (Guillaume & Rougemont, 2006) (under both the diallelic and continuous allele model) demonstrated that *r*_*G*_ and the correlation among effect of loci are different measures when genetic correlation *r*_*G*_ is selection-induced (Appendix A and Fig. S4). The individual-based simulation also suggested that the underlying genetic architecture of *r*_*G*_ may indeed have potentials to infer magnitude of mutational correlation in creating genetic correlations. Furthermore, such different mechanisms may be a determinant of genetic correlation stability (Appendix A and Fig. S5)

### Identifying loci with high degree of horizontal pleiotropy (HP) for yeast morphological traits

We next applied our approach in a yeast morphology dataset, where 374 recombinant strains of yeast cells were imaged for, on average, 800 fixed, stained cells per strain using high-throughput microscopy (Geiler-Samerotte *et al*., 2020). In total, measurements of 167 morphological traits were acquired. The patterns in this large dataset could offer a empirical picture of how the presence of HP may offer information about two mechanisms causing genetic correlation.

As described above, we define developmental bias *r*_*D*_ as the consistency of pleiotropy for a subset of variants where outliers (top HP loci) are removed. For example, the effect size distribution (exclude top HP loci) for two pair of traits are shown in Fig. 3. The red lines indicate the magnitude of *r*_*D*_ and the plot on the right is inferred to have a higher contribution from developmental bias. To identify outliers (top HP loci), we first calculate the correlation by individual-level product (Lea *et al*., 2019) for each trait pair across each locus. Outliers are then identified as the product falling outside 1.5 times the interquartile range (IQR) above the upper quartile and below the lower quartile of the distribution. We conducted robustness analyzes to show that changing the cut-off (1.5 IQR) does not qualitatively change the estimation of *r*_*D*_ (Fig. S6). In Fig. 3, these outliers are absent such that we can measure the innate correlation between two traits that results from developmental bias.

**Figure 3:**
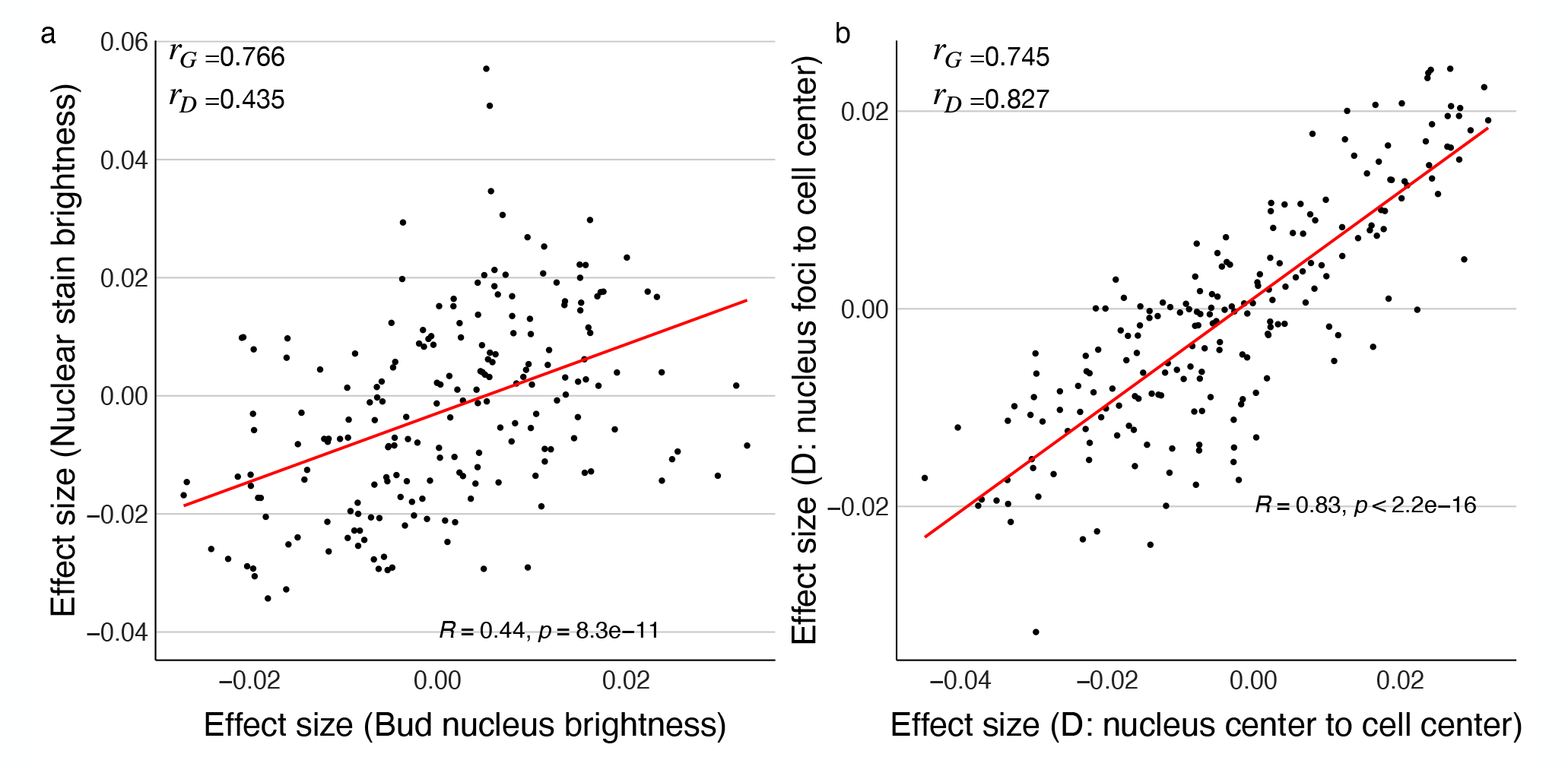
Empirical examples of the two scenarios illustrated in Fig. 1 demonstrating how estimated *r*_*D*_ can differentiate two trait pairs with similar *r*_*G*_ in yeast. These two trait pairs with similar strength of *r*_*G*_ (0.766 and 0.745) exhibit difference in *r*_*D*_ (0.435 and 0.827). **a**. The *r*_*G*_ between nuclear stain brightness and bud nucleus brightness is marginally higher than the **b**. *r*_*G*_ between nucleus foci-to-cell center and nucleus center-to-cell center. However, *r*_*D*_ between nuclear stain brightness and bud nucleus brightness is lower than *r*_*D*_ between nucleus foci to cell center and nucleus center to cell center (0.435 vs 0.827), suggesting a higher developmental bias and inherent inter-dependency between nucleus center-to-cell center and nucleus foci-to-cell center. Each point represents the additive effects for a single locus on each of two traits shown. Data are from (Geiler-Samerotte *et al*., 2020)

Effect size correlations, such as those depicted in Fig.3, can be affected by HP, but they also can be affected by LD. Thus, we performed LD pruning to subset the variants to remove highly correlated loci within the population (see Materials and Methods and Appendix B for general discussions of the LD effect). Therefore, in total, we present the effect size correlation against *r*_*G*_ in three settings: Default (using all genotyped variants), LD corrected, and outlier corrected (i.e., *r*_*D*_).

Fig. 4a presents the distribution of effect size correlations using all genotyped variants, only LD-pruned variants, or outlier-corrected variants (*r*_*D*_). LD does not exert effect on the patterns of effect size correlation in this dataset (blue and green lines in Fig. 4a are similar,two-sample Kolmogorov-Smirnov test, D = 0.017006, p-value = 0.3879), but horizontal pleiotropic loci, which we identified as outliers, appear to strengthen the effect size correlation (red line Fig. 4a shows that effect size correlations move closer to zero when outliers are removed). In principle, an outlier can either weaken or strengthen the effect size correlation. Yet, our results suggest a bias towards concordant effects between outliers and other loci, given that the distribution of effect size correlation is generally weaker under the outlier-corrected setting (Fig. 4a and b). In particular, points that deviate more from the unity line (*y* = *x*) may represent trait pairs that are more likely affected by horizontal pleiotropy (Fig. 4c).

**Figure 4:**
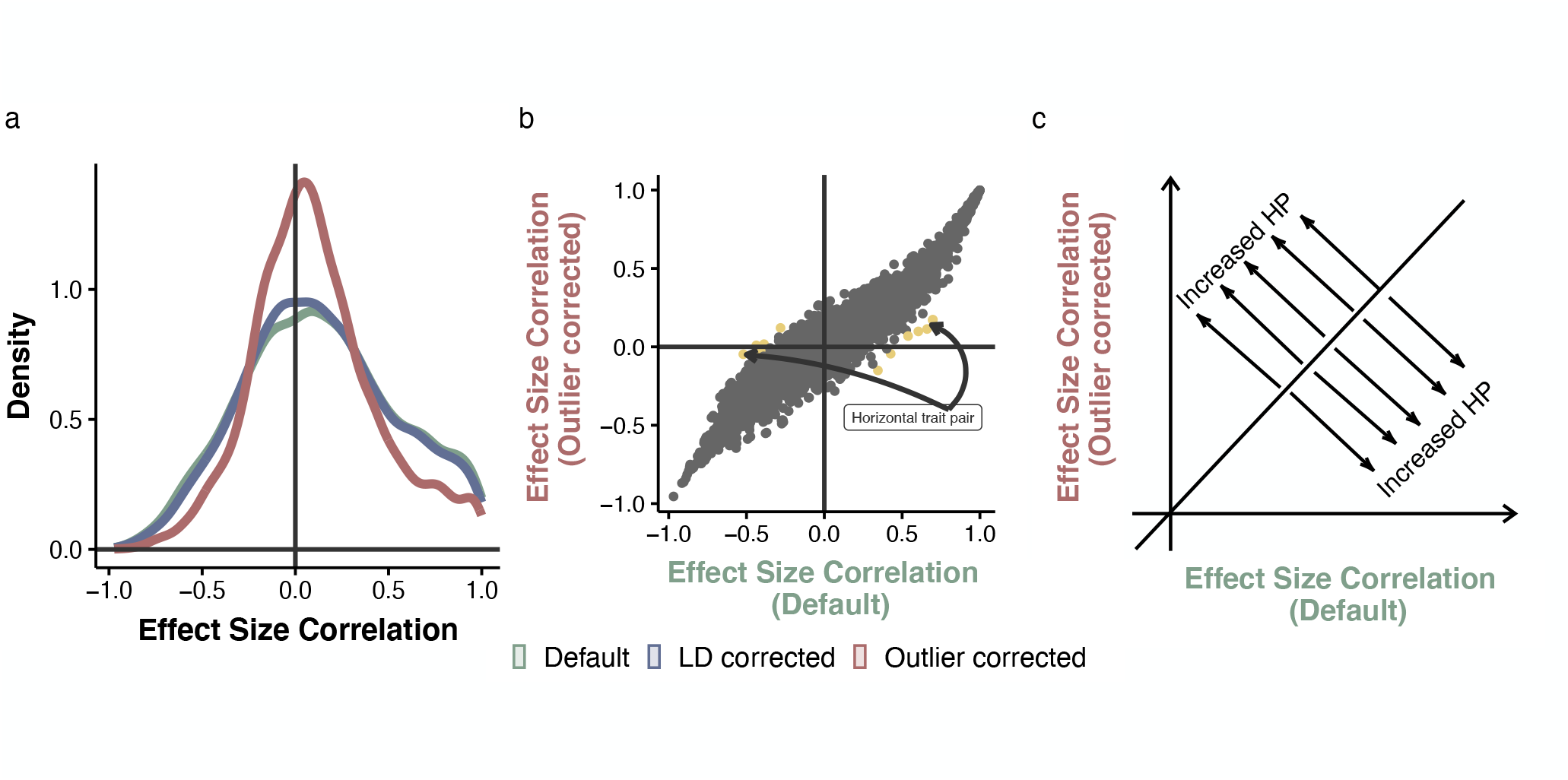
Re-analysis of 374 recombinant strains of yeast cells (Geiler-Samerotte *et al*., 2020) identifies horizontal pleiotropic trait pair. Each point in **b**. represents a trait pair from this empirical dataset. We consider three settings, summarized in **a**. and **b**.. **a**. Distribution of effect size correlations under three settings. Under the default setting, we included genome-wide markers to calculate correlations of effect sizes. LD pruned results include only the loci to the *r <* 0.5 within a chromosome. For outlier corrected effect size correlation, we excluded those outlier horizontal loci when calculating the correlation coefficient. **b**. The effect size correlation under default versus outlier corrected settings. Yellow dots denote trait pairs that are significantly affected by outliers correction (p-value *<*0.025). **c**. Conceptual figure showing where in the scatterplot **b** trait-trait effect size correlations have more contribution from horizontal pleiotropy.

Our method may also suggest which trait pairs are influenced by outlier loci, acting via horizontal pleiotropy. We identified those trait pairs significantly affected by the outliers (i.e., yellow dots in Fig. 4b). The outlier loci for these trait pairs are likely indicative of HP. Indeed, we confirmed that our method identifies two loci (L15.9 and L13.7, see Table 1) that presented the strongest evidence of HP in an earlier study (Geiler-Samerotte *et al*., 2020). Additionally, we found trait pairs with extremely high effect size correlations lacking evidence of horizontal pleiotropy (Fig. 4a, b); an example effect size distribution of a trait pair with exceptionally high effect size correlation – possibly reflecting a strong developmental bias between two traits – is shown in Fig. S7. In summary, we show that top HP loci may indeed affect *r*_*G*_ and that our method may be used to identify loci with high degree of HP and delineate *r*_*G*_. In addition, despite that there is no overall difference between the distribution of *r*_*D*_ and *r*_*G*_ (Fig. S3c), our approach can dissect *r*_*G*_ in empirical datasets as examplified in Fig. 3 that two trait pairs with similar *r*_*G*_ can be different in the magnitude of *r*_*D*_.

**Table 1:**
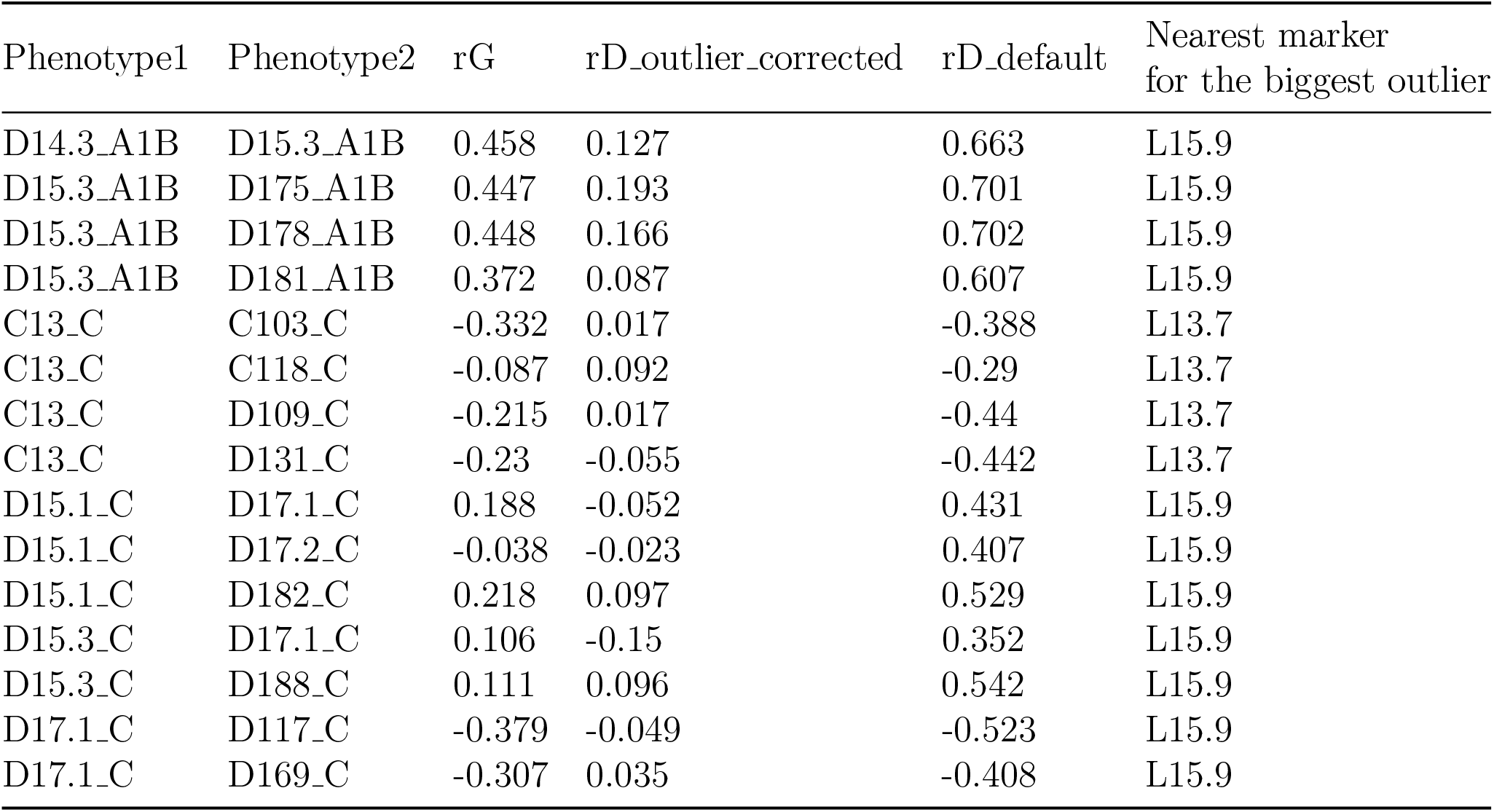
Details of horizontal trait pairs and nearest markers of driver loci.

### *r*_*D*_ predicts the stability of *r*_*G*_ following environmental perturbations

Genetic correlations between traits may alter the evolutionary trajectory of either trait (Schluter, 1996). Predicting the trajectory of trait evolution therefore may depend upon the stability of genetic correlations (Jones *et al*., 2003). Given the results of our individual-based models (Appendix A and Fig. S5), we reasoned that trait-trait correlations may be more stable if they exhibit higher locus-level correlation (presumably driven by developmental bias). Thus, we expected *r*_*D*_ to underlie the stability of *r*_*G*_ (Fig. 5a). To test whether our intuition is correct, we estimate *r*_*G*_ from a related yeast dataset, which describes correlations across yeast single-cell morphological features measured in three environments. Here, the environmental conditions are three concentrations of geldanamycin (GdA), a small-molecule inhibitor that binds the ATP-binding site of the chaperone Hsp90, thus rendering it unable to perform its cellular function. We plotted absolute *r*_*D*_ with changes of genetic correlation (*r*_*G*_) for each pair of traits at the three drug concentrations (Fig. 5). The results show that, as *r*_*D*_ increases, the changes of *r*_*G*_ decrease (Fig. 5 b).

**Figure 5:**
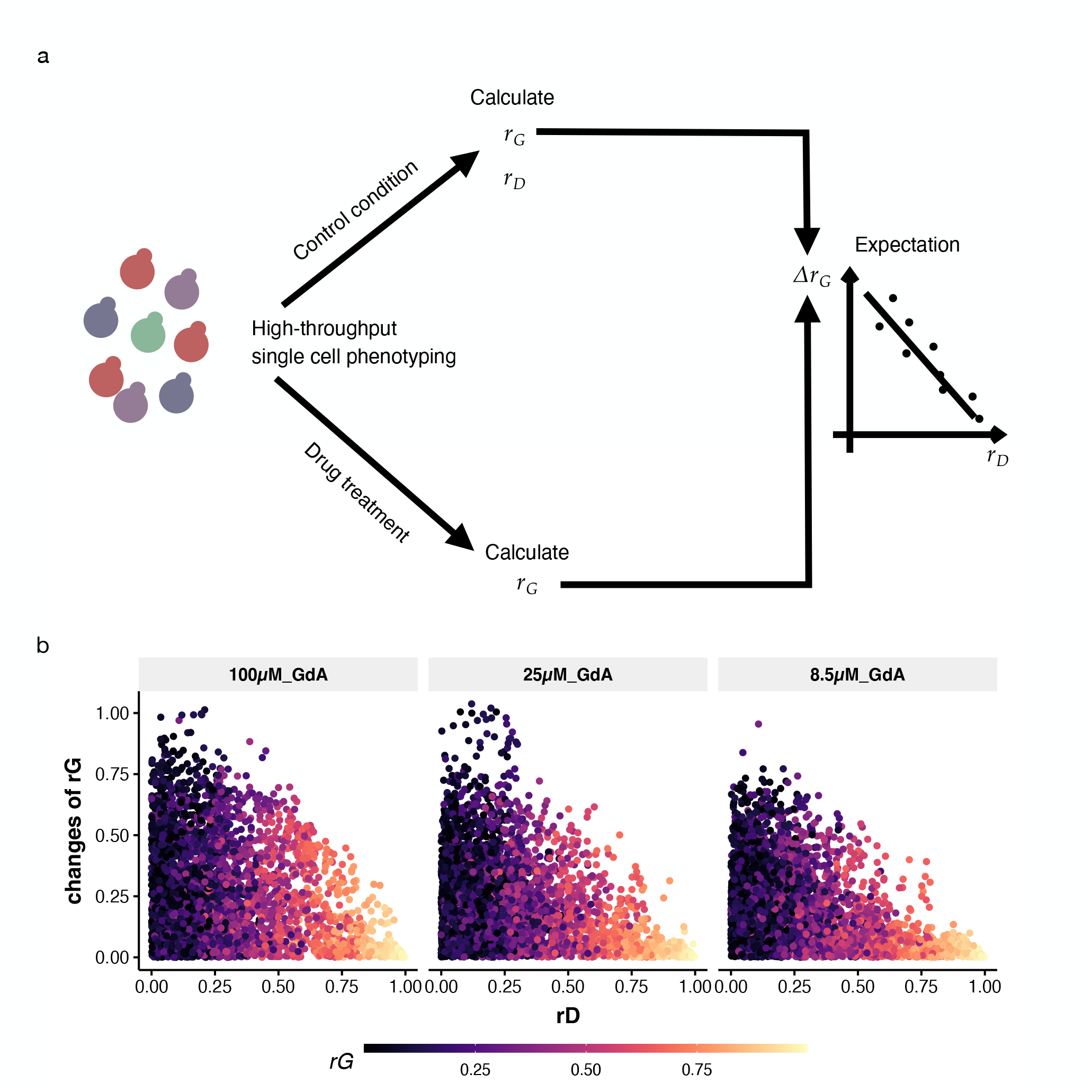
**a** Conceptual representation showing how high *r*_*D*_ could lead to stability of *r*_*G*_. Many morphological traits are measured in a yeast biparental population using single cell phenotyping (Geiler-Samerotte *et al*., 2020). These data are used to estimate *r*_*G*_ and *r*_*D*_ between traits. A subset of the yeast population is subjected to three levels of drug concentrations, representing three environmental conditions, and *r*_*G*_ is calculated for traits expressed in each of these treatments. We ask whether *r*_*D*_ is an indicator of Δ*r*_*G*_ by using a multivariable linear model: Δ*r*_*G*_ ∼ *r*_*D*_ + *r*_*G*_. We include *r*_*G*_ as a predictor because *r*_*G*_ itself can reflect its own plasticity. **b**. Scatter plots showing how *r*_*G*_ plasiticity in response to different drug concentration relates to *r*_*D*_. The color bar indicates *r*_*G*_ in control condition.

Since *r*_*G*_ is highly correlated with *r*_*D*_, to formally test whether under a given *r*_*G*_, *r*_*D*_ is informative in determining changes of *r*_*G*_ upon environmental perturbations, we conducted multivariable linear regression (Δ *r*_*G*_ ∼ *r*_*G*_ + *r*_*D*_, all variables are transformed to absolute value). The regression results (Fig. S8a) demonstrate that conditioning on *r*_*G*_ of a trait pair, *r*_*D*_ significantly negatively correlates with the changes of *r*_*G*_ across three drug concentrations. In other words, given a set of yeast morphology trait pairs with the same levels of *r*_*G*_, the magnitude changes of *r*_*G*_ would be smaller for trait pairs with larger *r*_*D*_, on average. Furthermore, we found that this effect of *r*_*D*_ is strongest under mild treatment perturbation (here, low concentration of geldanamycin) but becomes weaker as the drug concentration increases and, presumably, becomes more stressful to cells (Geiler-Samerotte *et al*., 2016). To account for the effect of collinearity between *r*_*D*_ and *r*_*G*_ (PCC = 0.939) on regression outcomes, we also report, here, null model simulated results (Fig. S8b and Fig. S9). Taken together, these results indicate that the estimated *r*_*D*_ may indeed contribute to the stability of *r*_*G*_ following environmental perturbations.

## Discussion

It has long been recognized that developmental integration is one cause of multivariate genetic constraint (Klingenberg, 2005; Pigliucci & Preston, 2004). On the other hand, genetic constraint can also reflect correlational selection. However, dissecting the underlying mechanism(s) causing genetic correlation is challenging. Here, we exploited data that have been overlooked to revisit the concept of pleiotropy and dissecting the mechanisms in producing genetic correlation. Assessing the consistency of pleiotropy by measuring the effect size correlation across many genomic loci provides a possible framework to explore the mechanisms of genetic correlation. The central messages from our analyses are three-fold.

First, consistency of pleiotropy inferred from genetic crosses provides an indicator of genetic correlation stability following environmental perturbations. Our stability analyses in empirical datasets (Fig. 5, Fig. S8, Fig. S9) suggest that the higher the consistency of pleiotropy, the more likely a genetic correlation between two traits remains stable across environmental conditions. In other words, higher developmental bias leads to a smaller response of the genetic correlation to environmental changes. This may provide further insight into the observations of context-dependencies of environmental effect on **G**-matrices (Wood & Brodie III, 2015), with certain trait pairs exhibiting more stability while others showing greater plasticity across conditions.

Second, Mendelian genetic architecture for a given trait pair can increase the contribution of horizontal pleiotropy (HP) to genetic correlation (Fig. 2). Under such a scenario, genetic correlation as a summary statistic can not fully reflect the complex genetic architecture underlying a genetic correlation. In fact, evidence of discrepancies of effect between genetic background and the major loci abounds (Albert *et al*., 2008; Hall *et al*., 2006; Scoville *et al*., 2009; Stinchcombe *et al*., 2009). For example, in *Mimulus*, a major QTL contributes a negative covariance between stigma–anther separation and pollen viability, which is antagonistic to the overall positive genetic covariance between these two traits (Scoville *et al*., 2009). Furthermore, previous work suggests that we might expect to see more changes of **G**-matrix during evolution if traits have an oligogenic genetic basis rather than aligning with the infinitesimal model (Barton & Turelli, 1987, 1989; Lande, 1979). For example, Lande (Lande, 1979) emphasized that trait means typically change much more rapidly than trait (co)variances. Yet, changes of (co)variances can be quite rapid if there are underlying loci with large contributions to (co)variation. Similarly, in our present work, we show that under a infinitesimal model, genetic correlation presumably arises from developmental bias (Fig. 2) which, as our stability tests suggest, might be more stable across conditions (Fig. 5).

Third, our method allows us to identify top HP loci without measuring phenotypes across clonal individuals or cells. There is a long-standing interest in identifying horizontal pleiotropy in nature (Verbanck *et al*., 2018; Jordan *et al*., 2019; Bowden *et al*., 2018). One motivation for doing so is that evolutionary theory predicts that natural selection should limit horizontal pleiotropy (HP) because, as the number of traits that a mutation influences increases, the probability of the mutation having a positive fitness effect decreases (Zhang & Wagner, 2013; Orr, 2000; Pavlicev & Wagner, 2012; McGuigan *et al*., 2014). However, identifying cases of HP is difficult because genetic correlations do not always indicate HP. By discovering a way to dissect genetic correlation, we have discovered a novel way to identify candidate loci that act via horizontal pleiotropy. Our method of identifying HP can be broadly useful because it does not require measuring the trait correlations that are present across clonal cells. Thus, while previous methods (Geiler-Samerotte *et al*., 2020) are mainly useful for organisms that propagated clonally or that can tolerate strong inbreeding, our method can be applied more broadly.

### Can recombinant mapping population characterize M matrix?

Numerous previous studies used mutation accumulation lines to estimate mutational matrices (**M** matrices) as a means to understand the influence of mutations on shaping genetic correlation (Dugand *et al*., 2021; Houle *et al*., 2017). Dugand *et al*. (2021) discovered a significant similarity between **G** and **M** matrices, suggesting that mutations directly shape **G**. On the other hand, mutational correlations using mutation accumulation lines were found to be overall stronger than genetic correlations between traits (Dugand *et al*., 2021), which is consistent with our findings using a genetic cross(Fig. S3a). In our results, effect size correlations using all variants overall are stronger than genetic correlations (Wilcoxon signed-rank test, p-value *<* 2.2e-16), with a median absolute value of 0.280 and 0.206, respectively. After removing top HP loci as outliers, there is no significant difference between the distribution of effect size correlations (*r*_*D*_) and the genetic correlation across trait pairs (Fig. S3c, Wilcoxon signed-rank test, p-value = 0.8875). This suggests that, hypothetically, top HP loci drive the pattern that mutational correlations are overall higher than genetic correlations between traits. This also naturally raises several questions related to mutation accumulation lines, recombinant mutation, and developmental bias: Firstly, do mutation accumulation lines reflect the propensity of developmental systems to vary – developmental bias? As Rohner & Berger 2023 pointed out, the mutation accumulation experiment also captures the heterogeneous mutation rates across the genome. The mutation rates have recently reported to be adaptive and uneven across the genome in a study (Monroe *et al*., 2022). Although the term “mutation bias” is inclusive of developmental bias, it goes beyond it as the mutation bias captures the limitation and bias in base pair changes and mutation rates across the genome (James *et al*., 2023). Secondly, to what extent does recombinant reflect the effect of mutation in a mutational accumulation experiment? There are now increasingly accessible resources available for recombinant mapping populations, such as the recently developed multiparent panels and advanced intercross lines (Kover *et al*., 2009; Gage *et al*., 2020), offering a promising avenue to investigate the respective roles of mutation and selection. In particular, a recent study has used genetic cross to provide neutral expectations of evolution (Fraser, 2020). Thirdly, to what extent do trait covariance patterns due to mutations or environmental perturbations reflect the developmental bias? A recent study used fluctuating asymmetry of the left and right sides of the same organism as a measure of developmental bias (Rohner & Berger, 2023). The left and right sides of the same organism share the same genome and macro-environment but only differ in their microenvironmental inputs. Therefore, the development may generate asymmetry (i.e., noise) in morphological traits. The authors showed that developmental bias quantified using such noise in the dipteran wing predicts its evolution on both short and long evolutionary timescales (Rohner & Berger, 2023), which suggests that those mild perturbations may generate phenotypic outcomes more representative of developmental bias.

### The extent of pleiotropy

Pleiotropy describes the phenomenon in which a gene or a mutation affects more than one phenotypic trait. The concept and nuance of pleiotropy has had a prominent role and broad implications on genetics, evolution, and medicine (Klingenberg, 2008; Stearns, 2010; Promislow, 2004; Williams, 2001; Barton, 1990; He & Zhang, 2006; Otto, 2004; Wagner & Zhang, 2011; Des Marais & Juenger, 2010; Geiler-Samerotte *et al*., 2020). Conceptually, many possible scenarios can result in a pleiotropic effect, including mediated pleiotropy (i.e., vertical pleiotropy), horizontal pleiotropy, and other spurious pleiotropy such as linkage (Wagner & Zhang 2011; Solovieff *et al*. 2013 and see Appendix B). An enduring debate about the genetic architecture of traits centers around how much pleiotropic natural systems are (Paaby & Rockman, 2013; Zhang & Wagner, 2013). A key challenge is whether the effect of a single locus on correlated traits can be counted as pleiotropic effects,Wagner & Zhang 2011; e.g., are the depth and the width of a bird beak two characters? Thus, ignoring trait correlations may bias the estimation of pleiotropy. One possible solution is to consider the effective number of traits by looking at the eigenvalue variance of the phenotypic correlation matrix (Wagner & Zhang, 2011; Pavlicev *et al*., 2009; Wagner *et al*., 2008): The more dispersed the eigenvalues, the more interdependency of the traits. However, this approach likely biases the interdependency estimations of the traits, especially in the presence of major pleiotropic effect loci. For example, as an extreme case, even a single pleiotropic locus alone can drive trait correlations in spite of the low consistency of pleiotropy (Fig. S1 and also see (Agrawal *et al*., 2001)). Hence, in this case, the effective number of traits calculated via the phenotypic correlation matrix will be overestimated simply because there is a major effect pleiotropic locus – this does not necessarily mean two traits are inherently interrelated. Similarly, the bias is present if there is a major antagonistic loci against overall correlation of effect size. Instead, our analyses demonstrated that the consistency of effect sizes may provide a more appropriate way to measure inherent trait correlation, and hence effective trait dimensions.

## Materials and Methods

Sophisticated tools in the field of quantitative genetics have been developed to identify genetic loci which statistically explain phenotypic variance in quantitative traits to regions of chromosomes, so-called quantitative trait loci (QTLs). One of the fundamental metrics of quantitative genetics is the additive effect of a QTL, which represents the change in the average phenotype produced by substituting one allele for another (Lynch *et al*., 1998; Falconer *et al*., 1996). To better illustrate what we could exploit through the additive effect distribution, bivariate effect size distribution under two scenarios are shown (Fig. 1a,b), where both of two pairs of traits are affected by a major pleiotropic locus. In contrast, the consistency of pleiotropic effect throughout genome is different. This illustrative example may be extreme, but it implies that only analyzing the summary statistics such as genetic correlation or statistically significant loci in a genetic association study may lose information behind the genetic architecture. Such hidden information could be valuable when assessing the strength of mutation and developmental bias *r*_*D*_ in generating *r*_*G*_: If developmental bias and mutation contribute more to correlation between two traits, these two traits should be correlated and less dependent on specific variant/loci. i.e., there is consistency of the pleiotropic effect across genetic background (Fig. 1a). On the other hand, if two traits are genetically correlated simply because of several major pleiotropic loci for a given population, those small loci can have inconsistent effect between traits (Fig. 1b). We term such consistency of pleiotropy as developmental bias *r*_*D*_ and those loci with effect deviated from overall trend throughout the genome as horizontal pleiotropic (HP) loci.

Our conceptualization of developmental bias is similar to the definition of vertical pleiotropy or mediated pleiotropy Geiler-Samerotte *et al*. (2020). Indeed, the high consistency of pleiotropic effect implies vertical or mediated pleiotropic nature of loci. Yet, we here define the developmental bias as a trait-level metric, whereas the vertical or mediated pleiotropy most often refers to the effects of variants. In vertical pleiotropy, the traits themselves are biologically related, such that a variant’s effect on trait A inevitably causes the effect on trait B. Likewise, HP is defined as a variant or mutation causing an effect on two traits that are otherwise independent. Another distinction between developmental bias and vertical pleiotropy is that vertical pleiotropy frequently refers to a part of causal cascade, as exemplified by low-density lipoprotein (LDL) cholesterol levels causing the risk of heart disease (Geiler-Samerotte *et al*., 2020). Developmental bias, on the other hand, depicts the correlational structure among traits since many traits (e.g., morphological traits) do not necessarily exhibit direct causal relationships.

### Numerical simulations

To simulate multiple pairs of traits within a population, first genotypes were simulated through the function *simulateGenotypes* in PhenotypeSimulator (Meyer & Birney, 2018) with 226 SNPs (mimicking the actual number of loci in an empirical dataset; Geiler-Samerotte *et al*. (2020)) and 500 individuals, where the allele frequencies are either sampled from 0.05, 0.1, 0.2, and 0.5 or constant value 0.3 for a mapping population. Empirical and theoretical studies support that the phenotypic effect size for fixed loci should be exponential, with many loci of small effect and a few loci of large effect (Bomblies & Peichel, 2022; Orr, 1998). Accordingly, for each pair of traits, the additive effect for each SNP is sampled from a bivariate exponential distribution (bivariate Laplace distribution) with *µ* = (0, 0) and 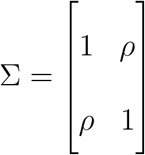. *ρ* is drawn from the uniform distribution (−1, 1). A shape parameter *γ* determines the distribution, where a smaller *γ* represents genetic architecture approximating one or a small number of large-effect loci (Mendelian genetic architecture, high kurtosis for effect size distributions) while a larger *γ* trends towards a polygenic infinitesimal model (low kurtosis for effect size distributions). We used *γ* of 1.0 and 0.5 in Fig. 2 left and right, respectively. Under horizontal pleiotropy scenario, *n* randomly selected SNPs (0 *< n <* 10) are forced to have horizontal pleiotropic effect (either concordant to or antagonistic with the rest of loci). The genetic correlation, *r*_*G*_, between traits *M* and *N* was calculated as the Pearson correlation 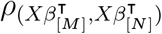, where *β*_[*M*]_ and *β*_[*N*]_ represents the effect size for trait *M* and trait *N* across genome, and 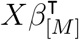 and 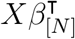 represent the genotypic values of trait *M* and trait *N*, respectively. *X* represents matrix consisting of genotypes for each individual. The developmental bias, *r*_*D*_, is calculated as the Pearson correlation coeffecient of effect size for traits *M* and *N*, where summations are taken over all loci except those assigned as horizontal pleiotropic SNPs (*n* loci):

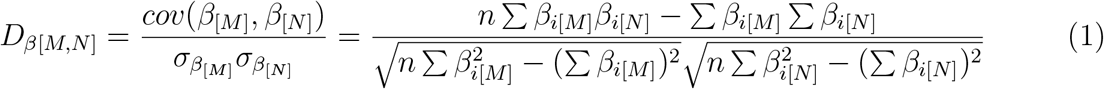

There exists considerable debate about regimes of allelic effect sizes and their effects on phenotypic evolution: in small steps, via changes of infinitesimally small effect, or in leaps via rare large effect loci (Orr, 2005). Additionally, classic work suggests that different genetic regimes may affect the rate of changes of genetic (co)variance (Barton & Turelli, 1987, 1989; Lande, 1979). Therefore, in addition to examining the effects of HP on the relationship between developmental bias *r*_*D*_ and genetic correlation *r*_*G*_, we performed these simulations under two genetic regimes, one with high polygenicity which causes low kurtosis in the distribution of effect sizes, and one with low polygenicity which causes high kurtosis in the distribution of effect sizes (Fig. 2a).

### Dataset retrieval and genetic correlations *r*_*G*_

The dataset comprises single cell morphology data for budding yeast *Saccharomyces cerevisiae* where, for each of 374 recombinant strains of yeast cells, approximately 800 fixed, stained cells were imaged using high-throughput microscopy (Geiler-Samerotte *et al*., 2020). A genetic linkage map of 225 loci typed in 374 recombinant segregants was previously developed (Gerke *et al*., 2009). The map covers the yeast genome at an average of 11cM intervals. 167 morphological features were estimated, including these representative examples: cell size, bud size, bud angle. Analysis of the original dataset assessed both genetic (between-strain) and environmental (within-strain) correlation using a multilevel correlation partitioning method (Bliese, 2013). The authors found that using this approach to estimate correlations has similar results as compared to a linear mixed model and variance component analysis.

### Horizontal pleiotropic loci identification and calculation of *r*_*D*_

If we ignore dominance, epistasis, and linkage disequilibrium, and assume two alleles per locus, the covariance components of a **G** matrix can be written as (Kelly, 2009):

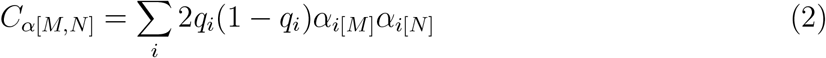

where *C*_*α* [*M*,*N*]_ is the additive genetic covariance between trait *M* and trait *N*. *q*_*i*_ is the frequency of first allele at loci *i* within a given population; *α*_*i*[*M*]_ and *α*_*i*[*N*]_ are the additive effects of that allele on trait *M* and *N*, respectively; summations are taken over all loci. Accordingly, a large effect QTL for trait *M* (high *α*_*i*[*M*]_) can make a minor contribution to the genetic covariance structure if allele frequency *q*_*i*_ is small.

We developed an approach to evaluate the horizontal pleiotropy (HP) and calculate *r*_*D*_ empirically. In brief, our method has three components: (a) detection of horizontal pleiotropic loci; (b) calculating *r*_*D*_ through effect size correlation excluding HP loci; (c) testing pairs with significant difference after outlier removal, identified as horizontal trait pairs.

We use the following procedures to identify HP loci: first calculate the normalized, demeaned, and element-wise product of outcome for each locus (Steiger, 1980):

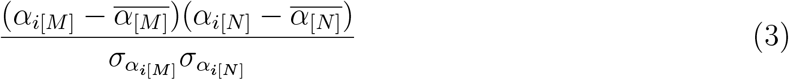

where 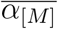 and 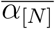 are the mean genome-wide additive effect size for trait *M* and *N*, respectively, and 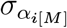 and 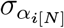 are standard deviations of additive effect size for trait *M* and *N*, respectively. The horizontal pleiotropic loci are defined as loci with the product falling outside 1.5 times the interquartile range above the upper quartile and below the lower quartile. The Pearson correlation coefficient is equal to the above average element-wise product of two measured traits. In other words, the outliers of these element-wise products represent the outliers when calculating the correlation, substantially deviating from the overall trend of bivariate effect size distribution.

Second, developmental bias (*r*_*D*_) is calculated as the Pearson correlation coefficient among the rest of loci written by:

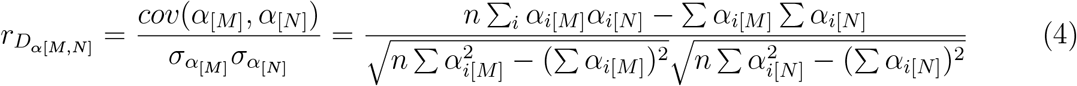

The additive effect size *α* estimated empirically from inbred line crosses and experimental mapping populations is calculated as: for the trait *M*, 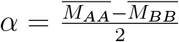 (Falconer *et al*., 1996), where *A* and *B* are two alleles of a locus. The summations of equation (4) are taken across all loci (with equal probabilities) excluding major horizontal pleiotropic loci.

Finally, to test the significant difference of effect size correlation before and after outlier removal, we use a cutoff of 1% FDR for all pairs of traits included in the yeast dataset. Those trait pairs significantly deviating from the mean are identified as the horizontal trait pair, indicating that horizontal pleiotropic loci may contribute to the genetic correlation *r*_*G*_.

## Data Availability

Data and code have been deposited in Github (https://github.com/haorancai/developmentalbias)

## Acknowledgements

The authors thank Matt Rockman, Luis-Miguel Chevin, and Tianzhu Xiong for helpful comments on an early draft. The authors thank John Kelly, Meghan Blumstein, and Jie yun for discussions.

